# Cryo-EM structure of the bacterial actin AlfA reveals unique assembly and ATP-binding interactions and the absence of a conserved subdomain

**DOI:** 10.1101/186080

**Authors:** Gulsima Usluer, Frank Dimaio, Shunkai Yang, Jesse M. Hansen, Jessica K. Polka, R. Dyche Mullins, Justin M. Kollman

## Abstract

Bacterial actins are an evolutionarily diverse family of ATP-dependent filaments built from protomers with a conserved structural fold. Actin-based segregation systems are encoded on many bacterial plasmids and function to partition plasmids into daughter cells. The bacterial actin AlfA segregates plasmids by a mechanism distinct from other partition systems, dependent on its unique dynamic properties. Here, we report the near-atomic resolution cryo-EM structure of the AlfA filament, which reveals a strikingly divergent filament architecture resulting from the loss of a subdomain conserved in all other actins and a novel mode of ATP binding. Its unusual assembly interfaces and nucleotide interactions provide insight into AlfA dynamics, and expand the range of evolutionary variation accessible to actin quaternary structure.

**Significance Statement:** Actin filaments are dynamic cytoskeletal elements that assemble upon ATP binding. Actin homologs are present in all domains of life, and all share a similar three-dimensional structure of the assembling subunit, but evolutionary changes to subunit have generated many different actin filament structures. The filament structure of the bacterial actin AlfA, which positions plasmids - small, circular DNA molecules that encode important genes - to ensure that each daughter cell receives a copy at cell division. AlfA is different from all other actins in two critical ways: it binds to ATP in a unique way, and it is missing a quarter of the conserved structural core. These differences explain unusual AlfA assembly dynamics that underlie its ability to move plasmids.

## Introduction

Actin is one of the most highly conserved eukaryotic proteins, with critical roles in processes as diverse as motility(1), cell shape(2, 3), organelle positioning(4) and cell division(5). Bacterial actins are involved in many of the same processes (6-8), and share evolutionarily conserved functional properties with eukaryotic actin: they form filaments whose assembly and disassembly is controlled by ATP binding and hydrolysis(9-14), their assembly dynamics are modulated by regulatory proteins(11, 15), and the filaments can serve as the basis for larger cellular structures(9, 16, 17). Actins all share a conserved structural core that has a complex topology of two domains (I and II), each subdivided into two subdomains (Ia, Ib, IIa, and IIb), with an ATP binding site between domains I and II(18). Five conserved sequence motifs (phosphate 1, connect 1, phosphate 2, adenosine, connect 2) in domains Ia and IIa surround the ATP binding site and have served to define members fo the family(19). The fold is also shared with Hsp70 and sugar kinases, which bind and hydrolyze ATP but do not form filaments. All members of the broader family undergo functionally important conformational changes upon ATP binding and hydrolysis that, in the actins, underlie assembly dynamics.

Despite these conserved features, bacterial actins exhibit far lower levels of sequence conservation than their eukaryotic counterpart. Unlike eukaryotic actin, where a single filament form has been adapted to multiple functions through a host of regulatory binding proteins, bacteria have evolved specialized actins for specific purposes that require fewer interaction partners. This has relaxed evolutionary constraints and allowed bacterial actins to explore a greater range of sequence diversity. The result is extensive divergence of bacterial actins at the sequence level, with corresponding variation in filament architecture, function, and dynamics.

A diverse subset of bacterial actins is involved in separation of plasmid DNA. Large, low-copy number plasmids often encode active segregation systems to ensure against stochastic loss when the host cell divides. Most segregation systems are composed of three elements encoded on the plasmid itself: a cytomotive filament to provide the force for plasmid movement, an adaptor protein that couples filament movement to the plasmid, and a centromere-like DNA region bound by the adaptor(20). Several different types of ATP-dependent cytomotive filaments have been adapted for plasmid segregation (21-23), with bacterial actins among the most widely distributed(12).

The most well studied actin-based segregation system is the *par* operon of the R1 multidrug resistance plasmid in *E. coli.* ParM filaments are dynamically unstable - they assemble upon ATP binding and hydrolyze ATP with kinetics that lag behind assembly, so that when the hydrolysis front reaches the end of a growing filament it catastrophically disassembles due to reduced stability of the ADP-bound state(10). ParM makes use of dynamic instability in a search and capture mechanism to segregate plasmids. Filaments nucleate spontaneously, and those that fail to encounter an adaptor-DNA complex eventually disassemble. When the ends of ParM filaments do encounter adaptor-DNA complexes their dynamic instability is suppressed, allowing processive growth that separates plasmids by pushing them toward opposite poles(15, 24).

The plasmid segregating actin AlfA, encoded by the *Bacillus subtilis* plasmid pLS32, was initially identified as an actin on the basis of the five conserved actin motifs, and like other actins AlfA forms ATP-dependent filaments both *in vivo* and *in vitro*(9, 13). Unlike ParM, however, AlfA is not dynamically unstable, forming stable filaments that remain assembled indefinitely in the ADP-bound state(9). Moreover, unlike ParM, which is structurally polar but grows at equal rates from both ends(10), AlfA filaments grow unidirectionally(25). AlfA filaments associate laterally into mixed polarity bundles, and extension of plasmid-bound filaments along bundles provides the mechanism of plasmid segregation. The adaptor protein AlfB regulates AlfA dynamics - free AlfB suppresses AlfA growth and promotes disassembly of ADP-bound filaments, while AlfB-DNA complexes nucleate AlfA filaments(25). These combined activities suppress spontaneous nucleation and ensure that filaments grow primarily from plasmids. Consistent with its unusual dynamics, our initial low-resolution structural studies of AlfA filaments revealed an unusual filament architecture, more ribbon-like and twisted than other actins(9). However, the relationship between this architecture and the dynamic properties of AlfA was unclear.

Here, we report the structure of AlfA filaments determined by cryo-EM at near-atomic resolution, which reveals the basis for its unique structural and functional characteristics. We show that AlfA lacks the canonical actin subdomain IIb, which plays important structural and functional roles in all other actins. AlfA polymerization interfaces have diverged extensively from other actins, and AlfA binds ATP through completely novel interactions with the adenosine base. These unique features of AlfA explain how it assembles stable filaments despite the loss of subdomain IIb, and why the filaments remain stable after ATP hydrolysis.

## Results and Discussion

### AlfA lacks subdomain IIb

In seeking clues to the unusual architecture from the AlfA sequence we carried out extensive sequence searches and multiple sequence alignment with bacterial actins. While AlfA was clearly identified as an actin on the basis of the five conserved actin motifs(13), sequence alignment of regions outside these motifs can be challenging due to the very low level of sequence identity. Beginning with alignments of only the closest relatives to AlfA and expanding the size of the sequence set, we were able to generate robust alignments showing AlfA is missing the canonical subdomain IIb (Fig. 1, Fig. S1). The closest homologs, with an average identity of ~20% to AlfA, constitute a family defined by the lack of IIb, consisting of actins primarily from gram-negative bacteria, and from several bacillus phages. The relatively limited number of actins in this family suggests that they have experienced a deletion of IIb during their evolutionary divergence from other actins.

**Figure 1.**
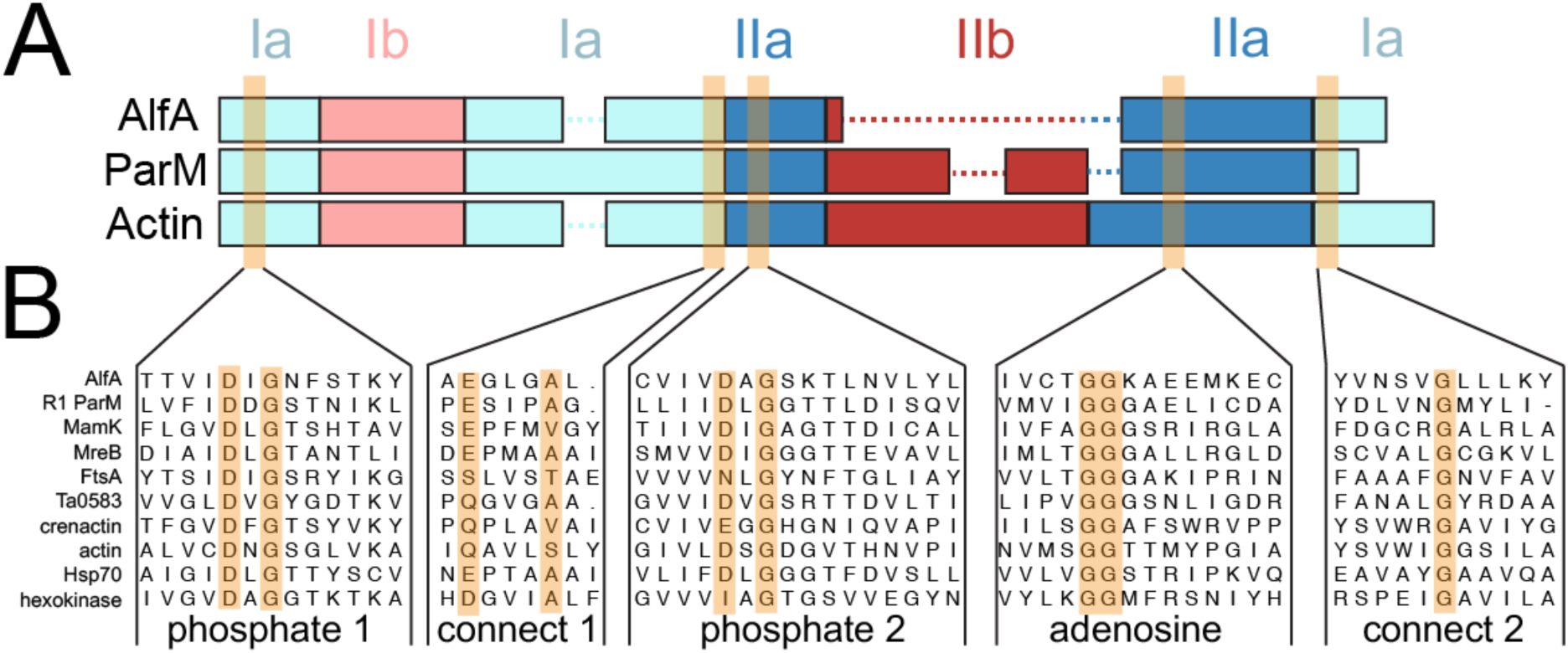
AlfA sequence conservation. A) Diagram of domain arrangements for AlfA, the bacterial actin ParM from the *E. coli* R1 plasmid, and vertebrate actin. The five conserved actin sequence motifs surrounding the actin binding cleft are highlighted in orange. B) Sequence alignments of AlfA with other actins, Hsp70 and hexokinase in the regions surrounding the five conserved motifs.

Subdomain IIb is critical for function in all other actins described to date. While IIb does not include any of the conserved ATP-binding motifs, it forms half of the binding pocket for the ATP adenosine base and contributes a significant fraction (17-35%) of the total surface area buried in filament assembly interfaces. In other actins ATP binding allosterically regulates polymerization by stabilizing a filament-bound ATP conformation whose major difference with the unbound conformation is the relative positioning of subdomains Ib and IIb (Fig. S2). The importance of IIb assembly interactions is highlighted by their role in ParM dynamic instability, where structural changes associated with ATP hydrolysis break IIb interactions with neighboring protomers, destabilizing the filament(26). The integral role of IIb in actin filament structure and function raises several questions: how can AlfA bind ATP with half of the canonical binding site missing, how is it able to form stable filaments without the IIb assembly interfaces, and how does ATP binding trigger polymerization? To answer these questions we determined the structure of the AlfA filament using cryo-EM.

### AlfA Filament Structure

At physiological salt concentrations AlfA filaments spontaneously assemble into bundles of variable thickness that are not well suited to high-resolution structure determination. However, bundles can be dissociated into single two-stranded filaments at high salt concentrations(9). We initially attempted to determine the cryo-EM structure of single AlfA filaments in 1M KCl, but background from high salt concentration limited the resolution of these reconstructions to about 12 Å. We then turned to a mutant we had previously designed, (two pairs of lysines, K21,K22 and K101,K102, mutated to alanines) that forms single filaments that do not bundle(25). These can be imaged at lower salt concentrations, making the sample better suited to high-resolution structure determination (Fig. S3).

We assembled the non-bundling AlfA mutant with the nonhydrolyzable ATP analog AMPPNP, and determined the structure of the filaments at 4.2 Å resolution by cryo-EM (Fig. 2, Fig. S4). The refined helical symmetry of the two-stranded filament was 157.7° rotation and 24.4 Å rise per subunit, giving a repeat distance of 394 Å (Fig. 2A). The two strands are parallel and offset by a half-subunit stagger. The repeat distance is considerably shorter than found in other actins, which range from 512 to 834 Å (27-32), yielding a highly twisted AlfA filament with only 8 subunits per turn of the two-start helix. The filament has a left-handed two-start twist, confirming our previous determination of handedness by tomographic reconstruction of negatively stained AlfA filaments (9).

**Figure 2.**
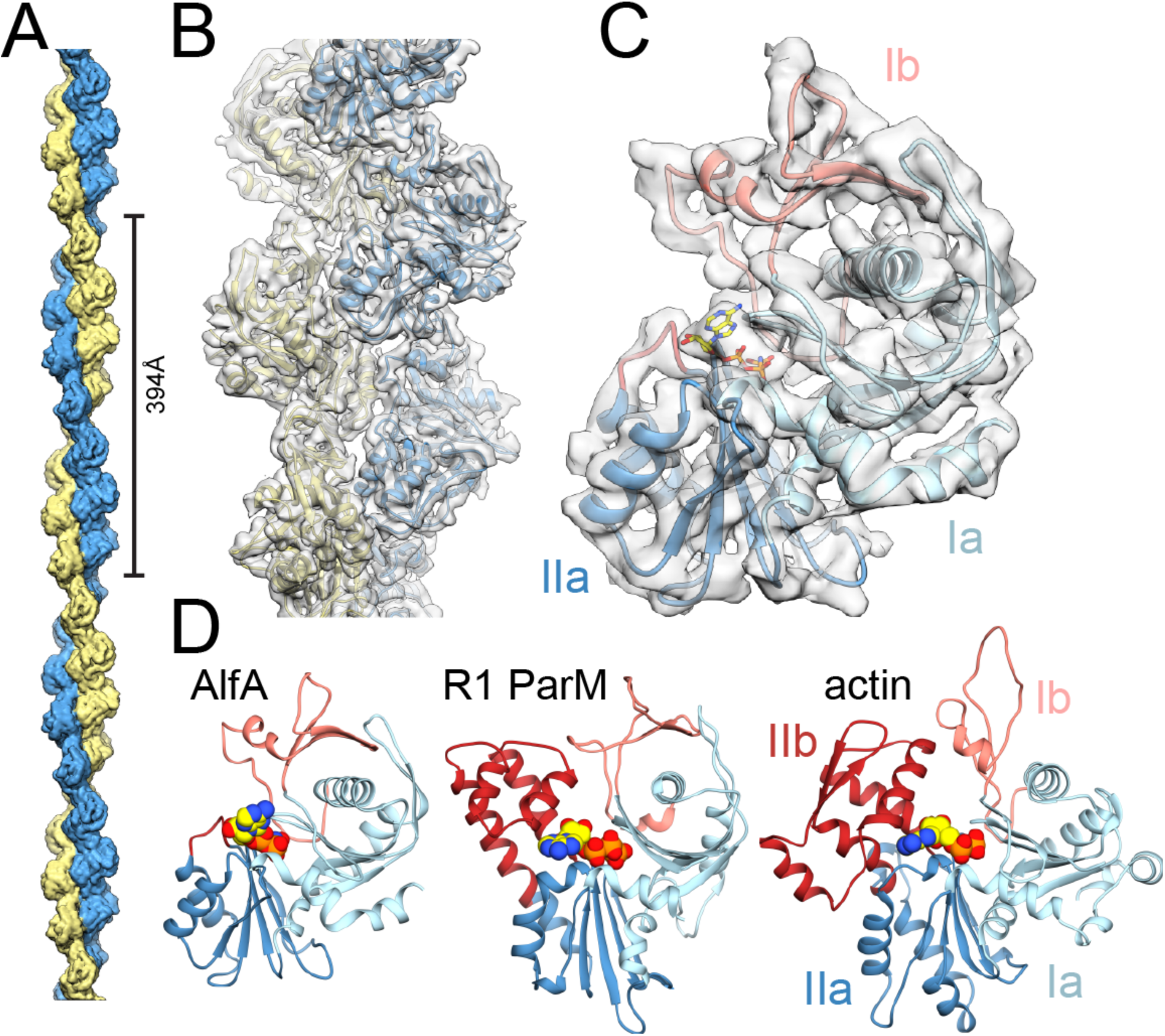
Cryo-EM structure of the AlfA filament. A) The two-stranded AlfA filament with a pitch of 394 Å. The pointed end is at top and barbed end at bottom, throughout all figures. B) The atomic model fit into a segment of cryo-EM density. C) A single AlfA protomer from the reconstruction colored by subdomain. E) Protomers of different actin filaments, with bound nucleotides in yellow, share a conserved fold, but AlfA is missing subdomain IIb.

The cryo-EM map has clearly defined secondary structure throughout, with bulky sidechains visible in some regions. We generated an atomic model using comparative modeling (33) followed by automated refinement (34), which covers the entire AlfA sequence and includes bound AMPPNP (Fig. 2 B,C; Fig. S4 C,D). Comparison of our atomic model with other actins confirmed that subdomain IIb is missing, replaced by a short five-residue loop (Fig. 2D). It is likely that the lack of IIb has reduced constraints on the helical symmetry of AlfA, making the more highly twisted architecture possible. The rest of the AlfA protomer has the typical actin fold, with subdomains Ia and IIa each built by a pair of alpha-helices packed against a five-stranded mixed polarity beta-sheet, and subdomain Ib consisting of a small three-stranded antiparallel beta-sheet and a short helix. Like ParM from the R1(35) and pSK41(36) plasmids, *Bacillus thurigensis* (37), and the archael actin Ta0853 (38), the subdomain Ib three-stranded beta-sheet of AlfA wraps around helix1 (residues 82-99), making contacts with a pair of anti-parallel beta-strands inserted after helix2 (residues 128-143) and burying most of helix1. This similarity suggests that this group of bacterial actins may share a more recent common evolutionary ancestor with each other than actins like MreB, MamK, crenactin, and eukaryotic actin that lack these features.

### A novel nucleotide binding mode in the AlfA filament

The density for AMPPNP is clearly defined in the cryo-EM map (Fig. S4). The overall backbone configuration of AlfA in the ATP binding region is conserved with existing actin structures, with average RMSD of 1.8 Å for backbone atoms between AlfA and protomers of other actin filaments. The three phosphates of AMPPNP are bound as in other actin structures, interacting with residues of the phosphate 1, connect 1, and phosphate 2 motifs. Strikingly, however, the adenosine base is rotated approximately 120° from the position it occupies in all other actin structures (Fig. 3A). Rather than packing against the adenosine motif in subdomain IIa, in AlfA the adenosine base is sandwiched between the phosphate 1 and connect 2 motifs in subdomain Ia. The ATP base stacks against the side-chain of Tyr255, while Phe12 packs against both the base and the ribose sugar (Fig. 3B). Both Phe12 and Tyr255 appear to be unique to AlfA (Fig. 3C). This novel ATP binding mode, with the base sandwiched between two parts of subdomain Ia, explains how AlfA binds ATP despite the lack of subdomain llb.

**Figure 3.**
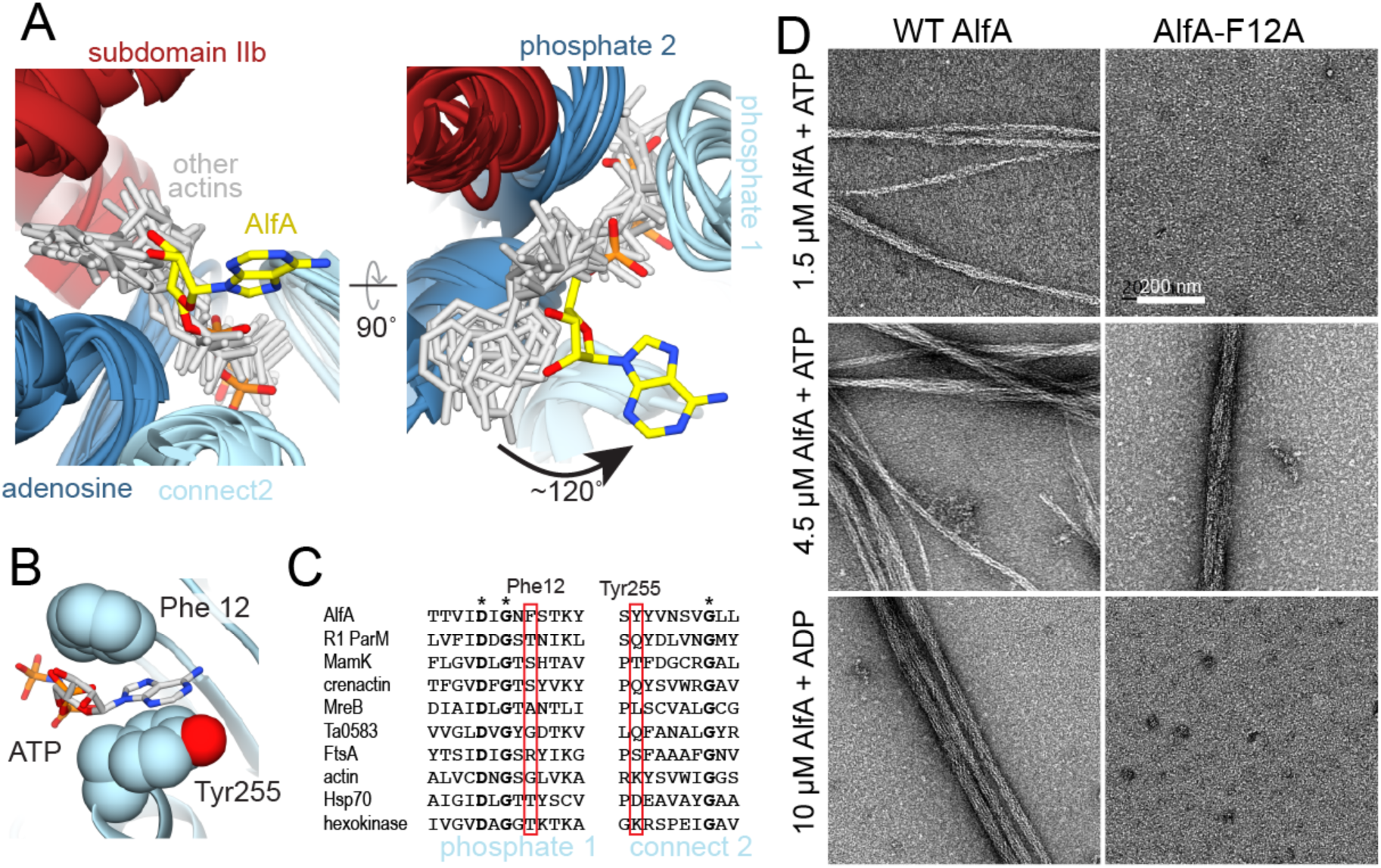
AlfA binds ATP through novel interactions. A) Structural alignment of ATP binding sites of AlfA and other homologs of the actin/Hsp70/sugar kinase family bound to nucleotide, with the proteins rendered as ribbons (colored by subdomain as in Figure 1), and nucleotides rendered as sticks (gray and yellow). The aligned structures are from cryo-EM filament reconstructions (PDB IDs actin:5JLF, MamK:5JLV, R1 ParM:5AEY, crenactin:5MW1), and crystal structures (PDB IDs MreB:4CZJ, Ta0583:2FSN, FtsA:1E4G, Hsp70:3KVG, hexokinase:2E2Q). Structural alignment was performed using just the regions around the conserved actin sequence motifs. B) In the AlfA filament structure the adenosine base is sandwiched between Phe12 and Tyr255 in subdomain Ia. C) Sequence alignment of the phosphate 1 and connect 2 actin motifs, with positions of Phe12 and Tyr255 highlighted in red. D) Negative stain electron micrographs of AlfA wildtype and F12A mutant in the presence of ATP and ADP. The F12A mutation is capable of assembling filaments but cannot maintain stable filaments in ADP. **[NEED image of high concentration, 100 or 200 uM]**

AlfA has only a four-fold difference in critical concentration between ATP and ADP(9), indicating that it does not distinguish strongly between di‐ and triphosphate nucleotides, and explaining why filaments remain assembled in the ADP-bound state. This is in stark contrast to ParM, which experiences dynamic instability and rapidly disassembles after ATP hydrolysis. We reasoned that the unique contacts AlfA makes with the ATP adenosine base provide the basis for its reduced discrimination between ATP and ADP. To test this, we generated a point mutant of Phe12 to alanine (AlfA-F12A) and tested its ability to assemble under different nucleotide conditions (Fig. 3D). The mutation increased the critical concentration for assembly to about 4 μM from the previously reported value of 2.5 μM (9). However, while wildtype AlfA assembles in ADP with a critical concentration of 10 μM, no assembly was observed with ADP for AlfA-F12A up to 200 μM protein. This highlights the importance of Phe12 in nucleotide binding and suggests that the unique ATP binding mode in AlfA is linked to the increased stability of AlfA filaments in ADP.

### AlfA assembly interactions

Actin filaments polymerize through two types of interface: head-to-tail longitudinal interactions that run along a single strand, and cross-strand lateral interactions. Longitudinal interactions define the structural polarity of actin filaments, with the two ends generally referred to as the barbed end (subdomains Ia and IIa) and the pointed end (subdomains Ib and IIb). In the AlfA filament longitudinal interactions bury about 1100 Å^2^ of surface area per protomer, while cross-strand contacts bury about 1400 Å^2^ (Fig. 4). Nearly all of the interaction surfaces are within subdomains Ib and IIa, with only very minor contributions from subdomain Ia, which plays a larger role in assembly of other actins (Fig. S5). While the total interface area per protomer is lower for AlfA than other actins, the fraction of its total surface involved in interfaces (20%) is comparable. However, the distribution of interfaces across the surface of AlfA is strikingly different from other actins. The lack of subdomain IIb means that AlfA longitudinal interfaces are less than half the size of the equivalent interfaces in other actins. However, AlfA has greatly extended cross-strand interfaces, which are 50-100% larger than those of other actins. The cross-strand interface is also mostly a single large patch running across subdomains Ib and IIa, compared to the distributed contacts scattered across all four subdomains in other actin filaments (Fig. 4A). The increase in the cross-strand interface area comes largely from interactions between a loop in subdomain Ib (residues 69-82) of one protomer with a helix in subdomain IIa (residues 201-211) that are brought into close contact due to the strong left-handed twist of the AlfA strands.

**Figure 4.**
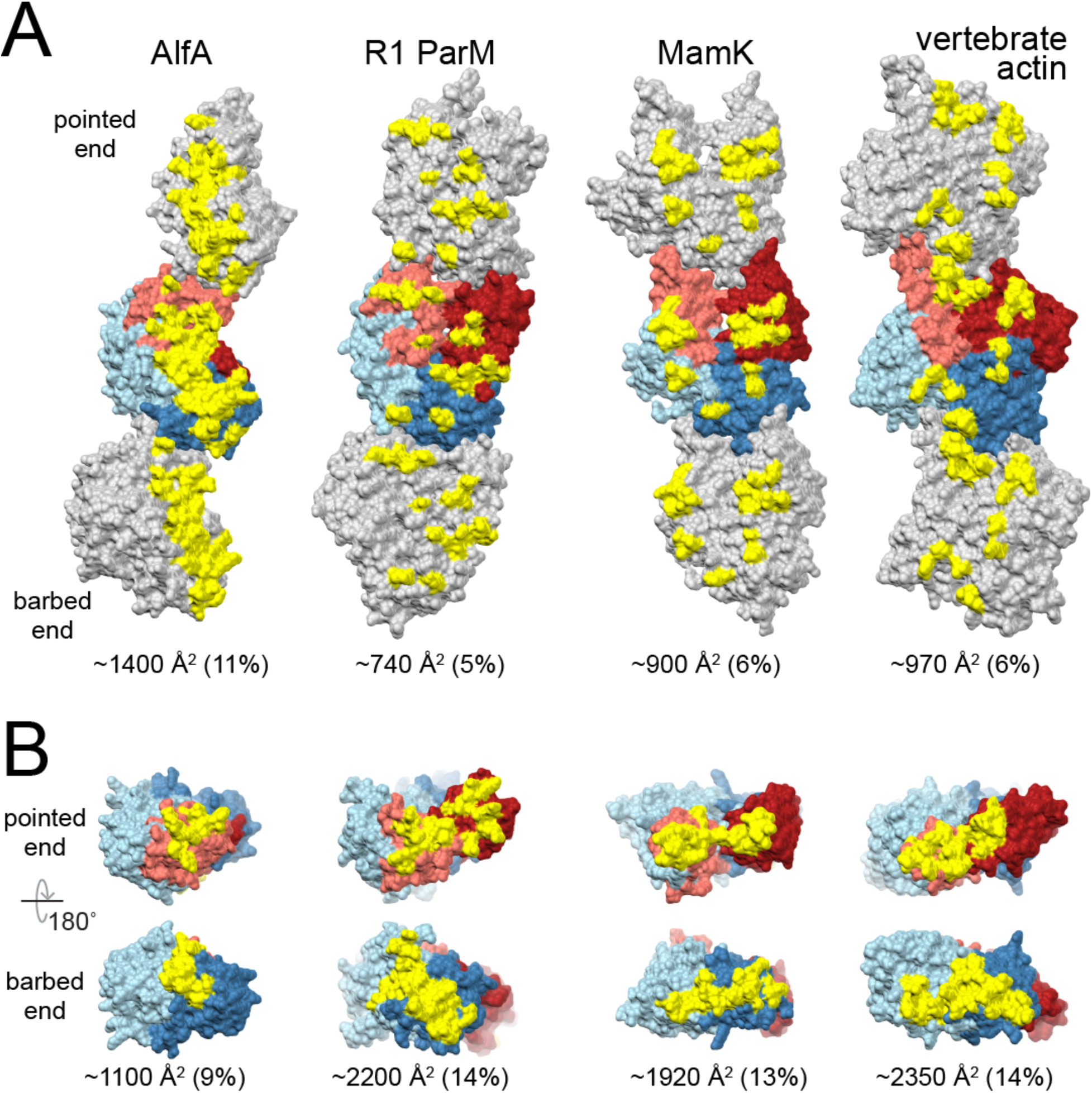
Increased AlfA inter-strand contacts compensate for missing subdomain IIb interactions. A) Three protomers from single strands of AlfA and three other actins are shown, with the central protomer colored as in Fig. 1. Residues involved in cross-strand interaction surfaces are colored yellow. B) Single protomers with residues involved in longitudinal interacting surfaces colored yellow. The size of each interface is given as both absolute area per protomer and as a fraction of the total surface area of the protomer.

We verified the AlfA assembly interfaces by generating mutations predicted to disrupt polymerization (Fig. S6). The mutations were designed at an early stage of cryo-EM structure determination, with a preliminary structure at ~12 Å resolution and a simple homology model of AlfA built by mapping the AlfA sequence onto the structure of R1 ParM. Of the six mutations tested, three prevented assembly and three failed to prevent assembly (Fig. S6C). Inspection of the mutation sites in the final high-resolution atomic model reveals that, in the cases where mutations fail to disrupt assembly the mutated residues are near interfaces but their sidechains are not in contact with other protomers (Fig. S6B). This indicates the accuracy of our atomic model of AlfA assembly interfaces.

### Consequences of AlfA filament architecture for dynamics and function

How ATP binding to AlfA promotes polymerization remains unclear from our AlfA filament structure. In other actins ATP-bound protomers are flattened in the filament relative to their free conformation, with the major change being a rotation between domains I and II around the connect 1 and connect 2 motifs. This conformational change results in large changes to the juxtaposition of subdomains Ib and IIb at the pointed end (Fig. S2). However, more subtle changes occur between Ia and IIa at the barbed end of the protomer, and assuming that AlfA undergoes a structural conversion similar to other actins these barbed end changes may be relevant to promoting filament assembly. The detailed nature of ATP-induced conformational changes in AlfA awaits a high-resolution structure of its apo conformation.

AlfA is unusual in that it exhibits extreme kinetic polarity, elongating almost entirely from one end, although which is the growing end has not been established (25). The structure of the filament provides a likely explanation for the asymmetry of subunit addition (Fig. 5). In other actin filaments assembly interactions involve both domain I and II whether adding at the barbed or pointed end. Interaction of both domains with the end of the filament stabilizes the flattened filament conformation of newly added protomers at either end. Similarly, at the AlfA barbed end the terminal protomer is bound through both subdomains Ib and IIa, which would create a conformationally stable new helical addition site. However, the terminal protomer at the pointed end is bound only via subdomain IIa. Being bound by only a single subdomain would potentially allow the terminal protomer to sample multiple conformational states and create a poorly defined, flexible helical addition site. This suggests that unidirectional growth of AlfA occurs at the barbed end, which would make it similar to actin, which grows more rapidly from the barbed than the pointed end, and distinct from ParM, which grows at indistinguishable rates from both ends (10).

**Figure 5.**
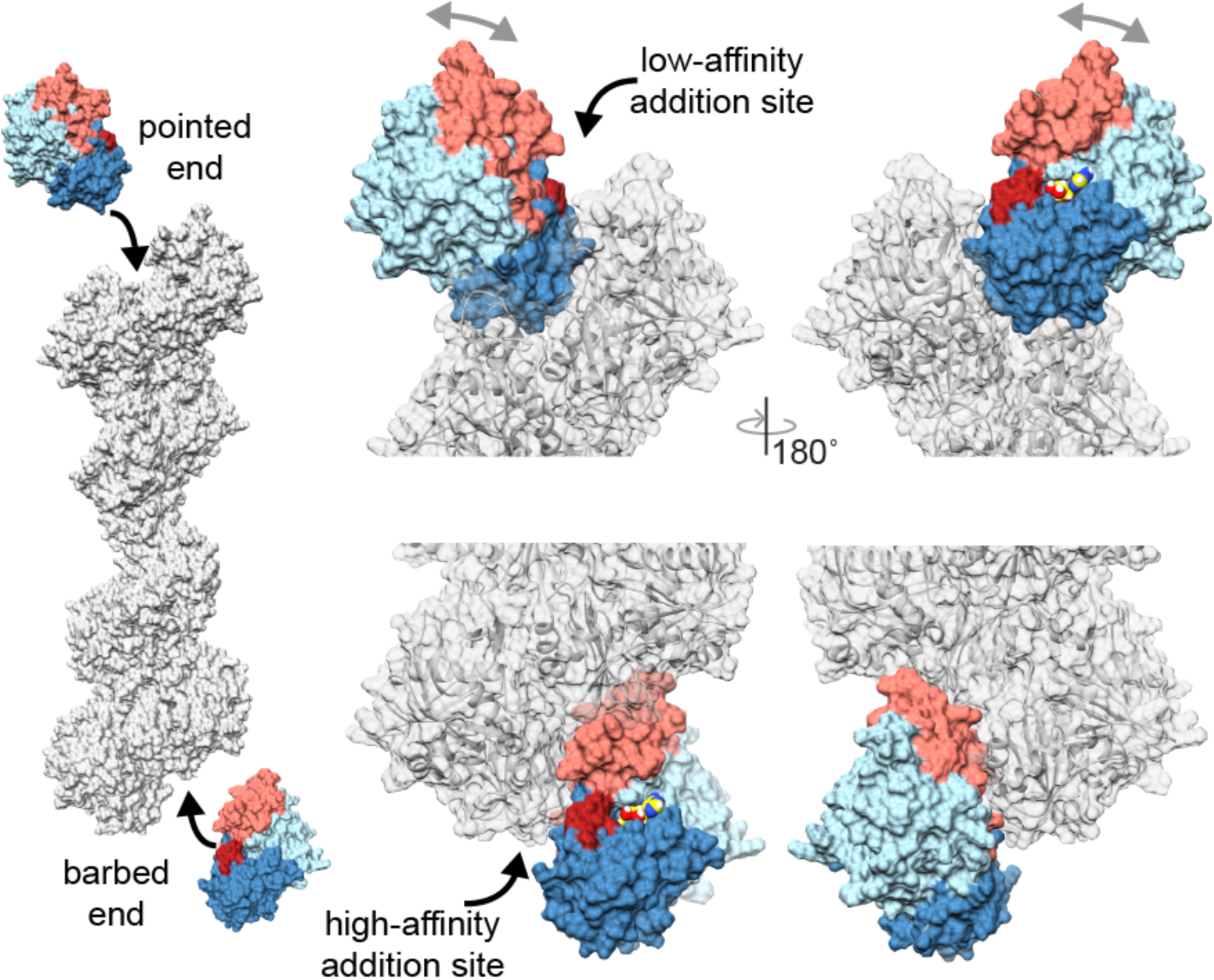
Structural differences between addition at barbed and pointed ends. New protomers would add to the pointed end only through interaction with subdomain IIa, which leaves domain I free to rotate relative to domain II (gray arrows). In contrast, addition at the barbed end involves interactions with both subdomains Ib and IIa, stabilizing a filament-bound conformation. Flexibility at the terminal pointed end protomer would create a very low affinity addition site, while the more rigid conformation of the terminal protomer at the barbed end would create a more defined high-affinity site, potentially explaining the observed unidirectional elongation of AlfA.

The adaptor protein AlfB regulates AlfA dynamics, nucleating AlfA filaments from plasmid DNA while simultaneously suppressing spontaneous nucleation. Moreover, after nucleation AlfB processively tracks the growing end of AlfA filaments, providing the basis for plasmid segregation (25). From the structural analysis of AlfA polarity above, suggesting filaments grow from the barbed end, we would also predict that AlfB binds at the barbed end. This would be similar to interaction between ParM and its adaptor ParR, which binds in a cavity between subdomains Ia and IIa that is only fully exposed at the barbed end. A similar situation may exist for AlfB binding to AlfA, which has a cavity in the region corresponding to the ParR binding site that partially overlaps with a longitudinal assembly interface. Such an interaction between AlfA and AlfB may be the key the dual role of AlfB in both suppressing (as free AlfB) and promoting (when bound to plasmid) AlfA polymerization; understanding the molecular mechanism of this activity requires further biophysical characterization of AlfA-AlfB interactions.

## Conclusions

The combination of altered domain architecture and novel mechanism of ATP binding give rise to the uniquely stable, highly twisted filament structure and unusual polymer dynamics of AlfA. The lack of a canonical actin subdomain is not unprecedented among bacterial actins, as FtsA, part of the cell division machinery, lacks subdomain Ib (39). However, in FtsA another domain is inserted at the barbed end of domain Ia, which makes contacts that compensate for the lost interaction surfaces(40). AlfA, on the other hand, has compensated through altered assembly interfaces, including a more extensive and continuous cross-strand interface. The divergent structure of AlfA highlights the extreme evolutionary plasticity of actin filament quaternary structure. This property has been exploited by bacteria to generate a broad range of actin filaments with unique dynamic and functional properties tuned to a wide variety of specific cellular functions. Given that only a small number of bacterial actins have been structurally and functionally characterized it is likely that further functionally important variation in filament morphology and dynamics remains to be discovered.

## Methods

### Sequence comparison

The sequence for AlfA was subjected to BLAST search (41), yielding only three matches of significant homology. These three matches were subjected to BLAST, increasing the size of the pool of AlfA homologs missing subdomain IIb, and this processes was repeated until no new homologs with the AlfA domain architecture were found. Multiple sequence alignment were calculated with MAFFT (42), using AlfA and the pool of close homologs and large representative samples of other bacterial actin familes (MamK, MreB, ParM, Alp12, Alp7).

### AlfA expression constructs

Previously described untagged expression constructs using a codon-optimized *alfa* gene were used to express wildtype (pJKP100) and non-bundling (pJKP102) AlfA(9, 25). AlfA-F12A was generated by site-directed mutagenesis of pJKP100. For non-assembling mutants the AlfA coding region was cloned into pSMT3-Kan (43), which inserted a His-SMT3/SUMO tag at the N-terminus of AlfA. The tag can be cleaved by ULP1 protease, leaving only two residual non-native residues at the N-terminus.

### Protein expression and purification

Recombinant AlfA constructs were expressed from IPTG inducible promoters in *E. coli* C43 cells at 18 °C overnight, as previously described (9). Wildtype, non-bundling AlfA, and AlfA-F12A were purified using a protocol similar to previous studies (9, 25), using cycling between polymerized and unpolymerized states as an initial bulk purification step. Cell pellets were resuspended in lysis buffer (25 mM Tris pH 7.5, 300 mM KCl), lysed by sonication, and the lysate cleared by ultracentrifugation for one hour at 4 °C at 34,000 rpm in a Type 50.2 Ti rotor (Beckman Coulter). AlfA was polymerized in the cleared lysate by addition of 5 mM ATP and 12 mM MgCl_2_ on ice for 30 minutes, and then pelleted by ultracentrifugation for one hour. The supernatant was discarded, and pelleted filaments were resuspended in depolymerization buffer (25 mM Tris pH 7.5, 300 mM KCl, 5 mM EDTA) then dialyzed overnight against the same buffer. Unpolymerized material was removed by ultracentrifugation, and the polymerization-depolymerization cycle was repeated. The final soluble AlfA sample was then applied to a Superdex 200 size exclusion column in polymerization buffer, peak fractions were pooled and concentrated to 100-200 μM, then flash frozen in liquid nitrogen and stored at -80 °C or stored for up to a week at 4 °C.

Cycling between polymerized and unpolymerized states could not be used to purify mutants designed to interfere with assembly. Instead, these were purified by Ni-NTA affinity chromatography, the His-SMT3/SUMO tag removed by cleavage with ULP1, followed size exclusion chromatography on a Superde× 200 column in polymerization buffer. Polymerization of wildtype AlfA purified in the same way from the same expression vector was indistinguishable from polymerization of untagged wildtype AlfA.

### Negative stain electron microscopy

Wild-type and mutant AlfA samples were polymerized for 15 minutes at room temperature in polymerization buffer plus 1 mM nucleotide and 1 mM MgCl_2_. Samples were applied to glow-discharged 400-mesh carbon-coated grids, and negatively stained with 0.7% uranyl formate (44). Images were obtained on an FEI Morgagni microscope operating at 100 kV, at 22,000x magnification, recorded on an Orious CCD camera (Gatan).

### Cryo-EM data acquisition

Non-bundling AlfA was assembled at room temperature, at 5 μM AlfA in polymerization buffer with 5 mM AMPPNP and 5 mM MgCl_2_ added. Samples were applied to glow-discharged C-FLAT 1.2/1.3-4C holey carbon grids (Protochips, Inc.) and plunge frozen in liquid ethane in a Vitrobot Mark IV vitrification device (FEI, Co.). Data were collected with an FEI TItan Krios microscope operated at 300 kV on a K2 Summit direct electron detector (Gatan, Inc.) in operating in super-resolution mode with a pixel size of 0.5 Å/pixel. Movies were recorded for 7.2 s, with 0.2 s frames, 72 e^-^/Å^2^ total dose per movie. Leginon was used for automated data acquisition (45).

### Cryo-EM image processing

Movies were aligned, dose-weighted, and Fourier binned using MotionCor2 (46). Defocus parameters were determined from the unweighted aligned sums using GCTF (47). Filaments were automatically identified using Relion (48), and extracted in overlapping 448 Å boxes using a step size of 25 Å to match the AlfA helical rise. This yielded 123,296 boxed segments. Helical segments were subjected to reference-free two-dimensional classification in Relion, and poorly aligning segments were rejected from further processing, leaving a data set of 113,222 segments for three-dimensional processing.

An initial reconstruction was calculated using iterative helical real space reconstruction in SPIDER, essentially as described (30, 49, 50). This model was low-pass filtered at 60 Å and used as an initial model for helical refinement in Relion (48, 51, 52). After initial gold-standard helical refinement using a spherical mask yielded a structure at about 5 Å resolution, helical segments were subjected to the Relion particle polishing routine, and refinement was continued using a shape-based soft-edged mask enclosing ~6 AlfA protomers (Supplementary Fig. 4B). The final reported resolution is from a Fourier shell correlation curve corrected for masking artifacts.

### Atomic model building and analysis

An initial structure of the AlfA protomer was generated asymmetrically using RosettaCM (33) for comparative modeling into the EM density map, using a diverse set of actin atomic structures as templates (PDB IDs: 1JCE, 2FSJ, 2ZGY, 3I33, 3JS6, 4APW, 4B1Y, 4KBO, 4PL7, 4RTF, 4XE7, 4XHP, 5EC0, 5F0X, 5LJW). Several loop regions (residues 65-83, 36-44, 195-215) were then rebuilt in Rosetta in the context of the helical lattice. This was followed by automated refinement of the entire structure in Rosetta using helical symmetry constraints using the protocol described by Wang, et al. (34). Finally, some sidechain rotamers were adjusted manually to improve fit to density.

The sizes of interacting surfaces between domains in AlfA and other actin filaments were calculated using the PDBePISA server (53). All cryo-EM structures and atomic models were visualized and figures prepared in Chimera (54).

## Acknowledgements

We are grateful to Daniel Southworth and the Life Sciences Institute at the University of Michigan for access to their microscopy facility, and to Julien Bergergon for helpful discussions. This work was supported by the Canadian Institutes for Health Research (298465 to J.M.K.), the Human Frontier Science Program (RGY0076 to J.M.K.), and The Scientific and Technological Research Council of Turkey 2213 Graduate Scholarship Program for Studies Abroad (to G.D.U.).

---

**Supplementary Figure 1.**
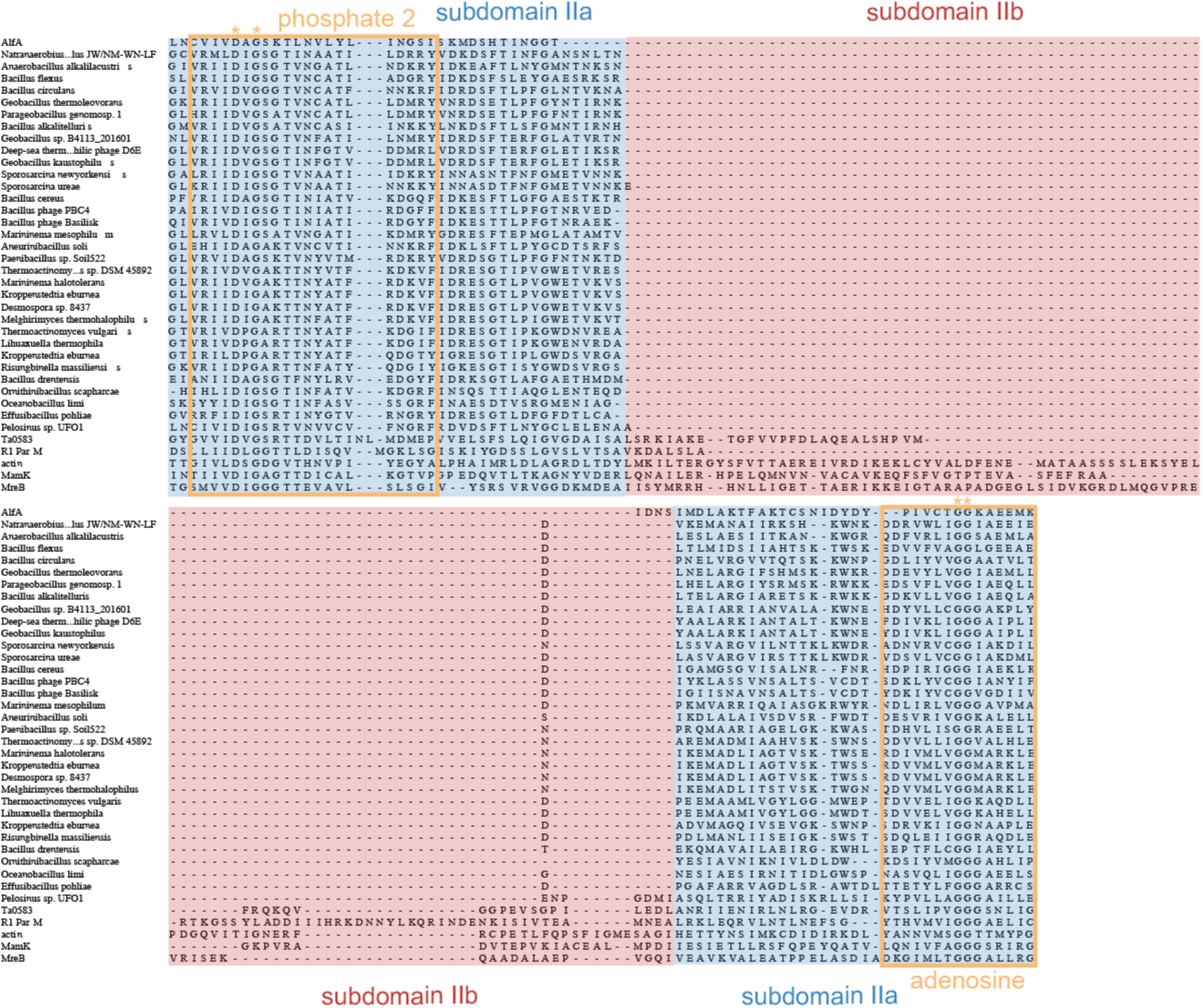
Sequence alignments of domain II. Multiple sequence alignment of subdomain IIb (red) and flanking regions in subdomain IIa (blue), demonstrates a family of bacterial and phage actins that lack subdomain IIb. The conserved phosphate 2 and adenosine motifs are outlined in orange and universally conserved residues highlighted with asterisks. The overall sequence identity between AlfA and other actins missing subdomain IIb is ~20%, while the identity to other bacterial actins is between 11% and 15%.

**Supplementary Figure 2.**
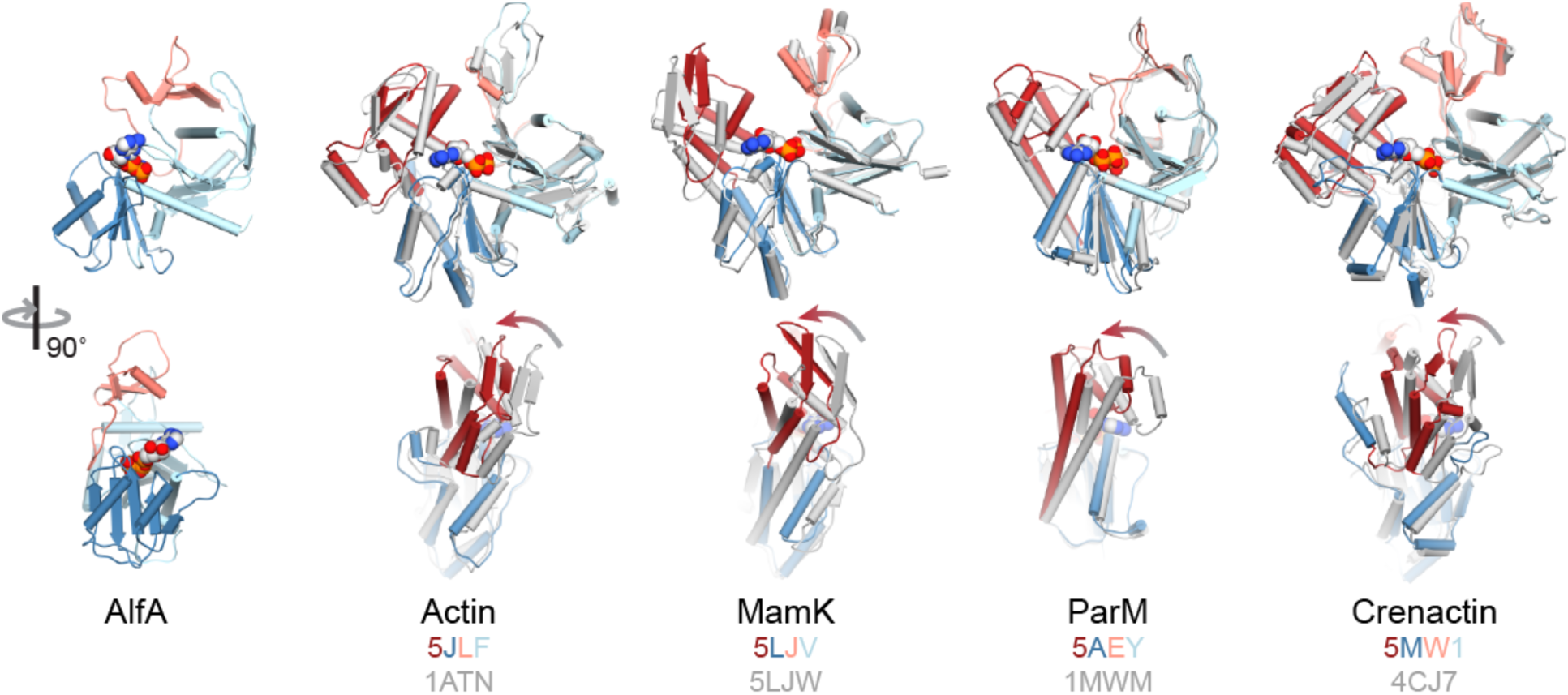
Conformational differences between free and filament-bound actins. Four different actins are show in their free conformations from crystal structures (gray) and filament-bound conformations from high-resolution cryo-EM reconstructions (color). In each case the major conformational change is a rotation of domains I and II relative to each other, yielding a flatter protomer in the filament. Arrows indicate the direction of the conformational change. The structure pairs were aligned on domain Ia in each case. PDB IDs are indicated for both states.

**Supplementary Figure 3.**
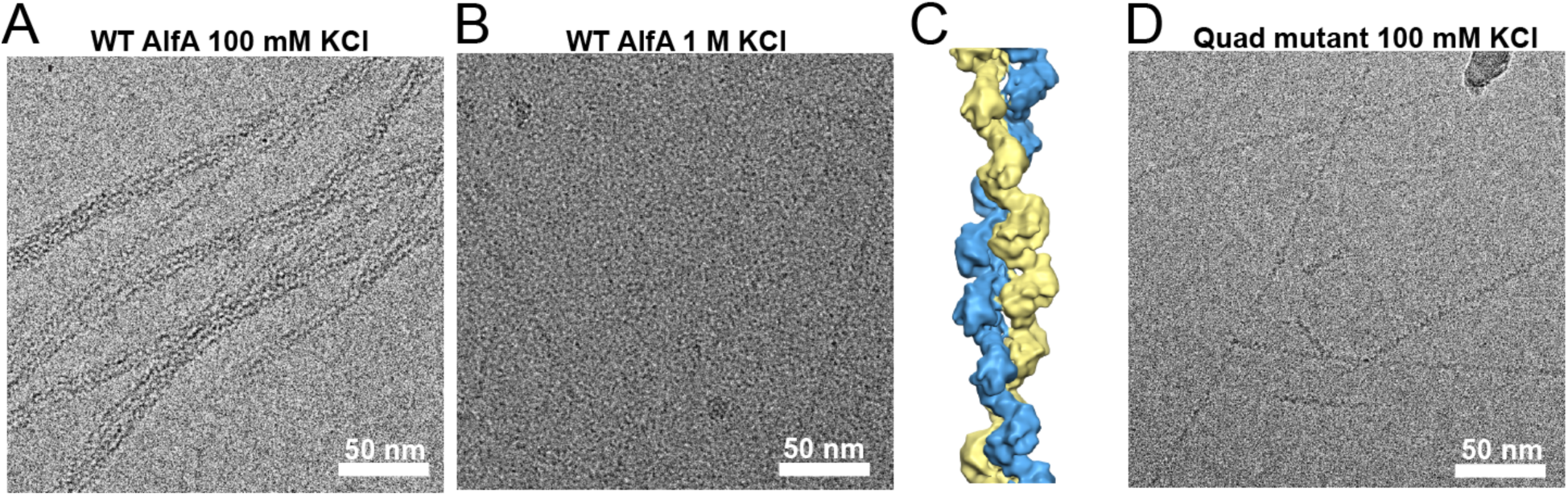
Optimization of AlfA cryo-EM samples. A) Cryo-EM image of AlfA at 100 mM KCl, where single filaments aggregate laterally into bundles with irregular thickness. B) Cryo-EM image of AlfA at 1M KCl, where bundle formation is inhibited but increased solvent density reduces contrast with the filaments. C) Reconstruction at 12 Å resolution of AlfA filaments in 1M KCl. D) Cryo-EM image of AlfA with four surface lysines (K21, K22, K101, K102) mutated to alanine (the ‘quad’ mutant), which inhibits bundling at low salt concentration. Images in B and D are both at -1.5 μm defocus.

**Supplementary Figure 4.**
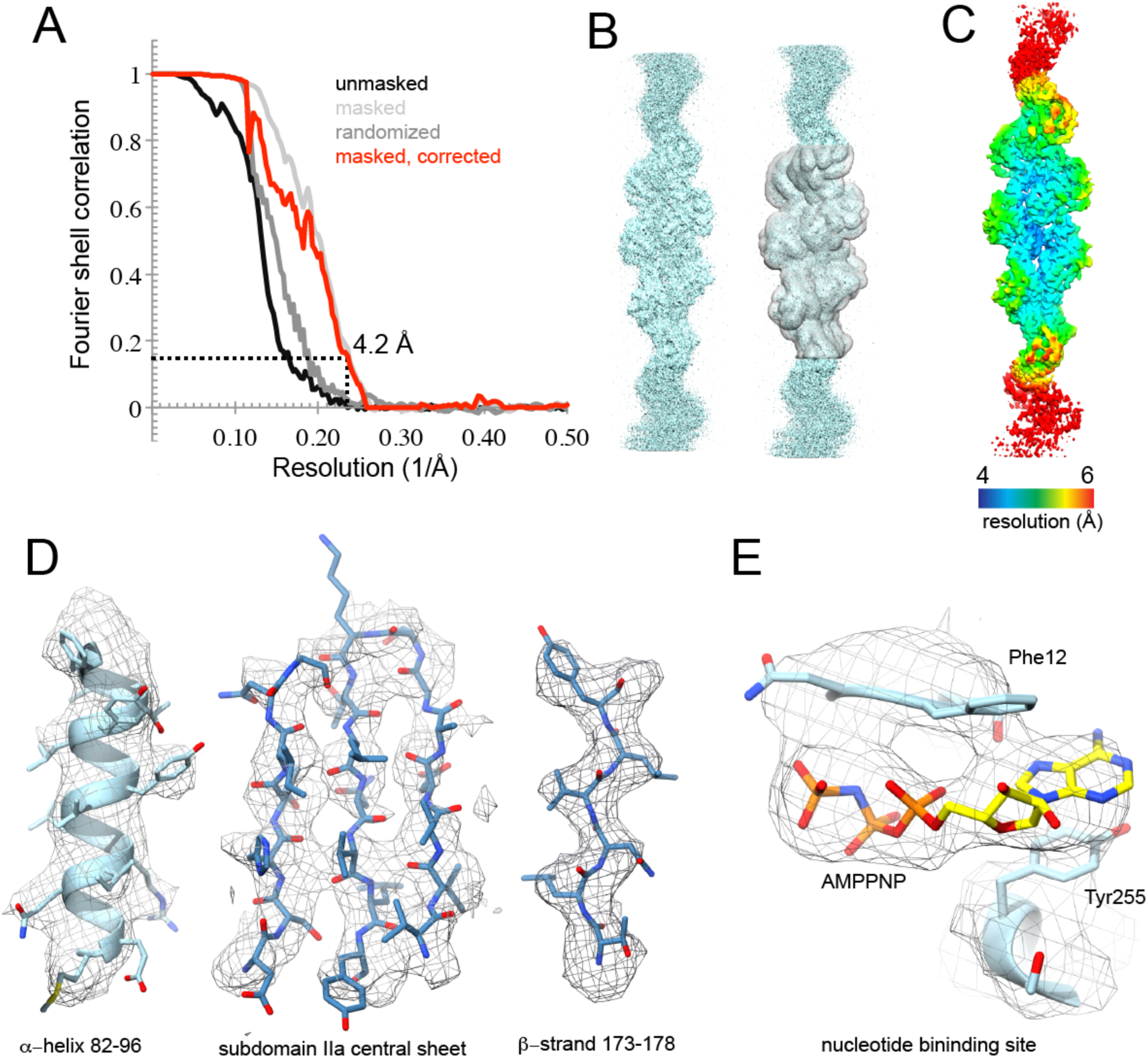
Cryo-EM Reconstruction of AlfA. A) FSC curves for final AlfA reconstruction. The final resolution calculated from the masked filament and corrected for masking effects is 4.2 Å. B) One unfiltered half map from the final reconstruction, shown with the mask used for calculating the FSC curves in (A) (right). C) Local resolution estimate calculated in Relion. D) Regions of representative density in the structure. E) Cryo-EM density in the nucleotide binding site.

**Supplementary Figure 5.**
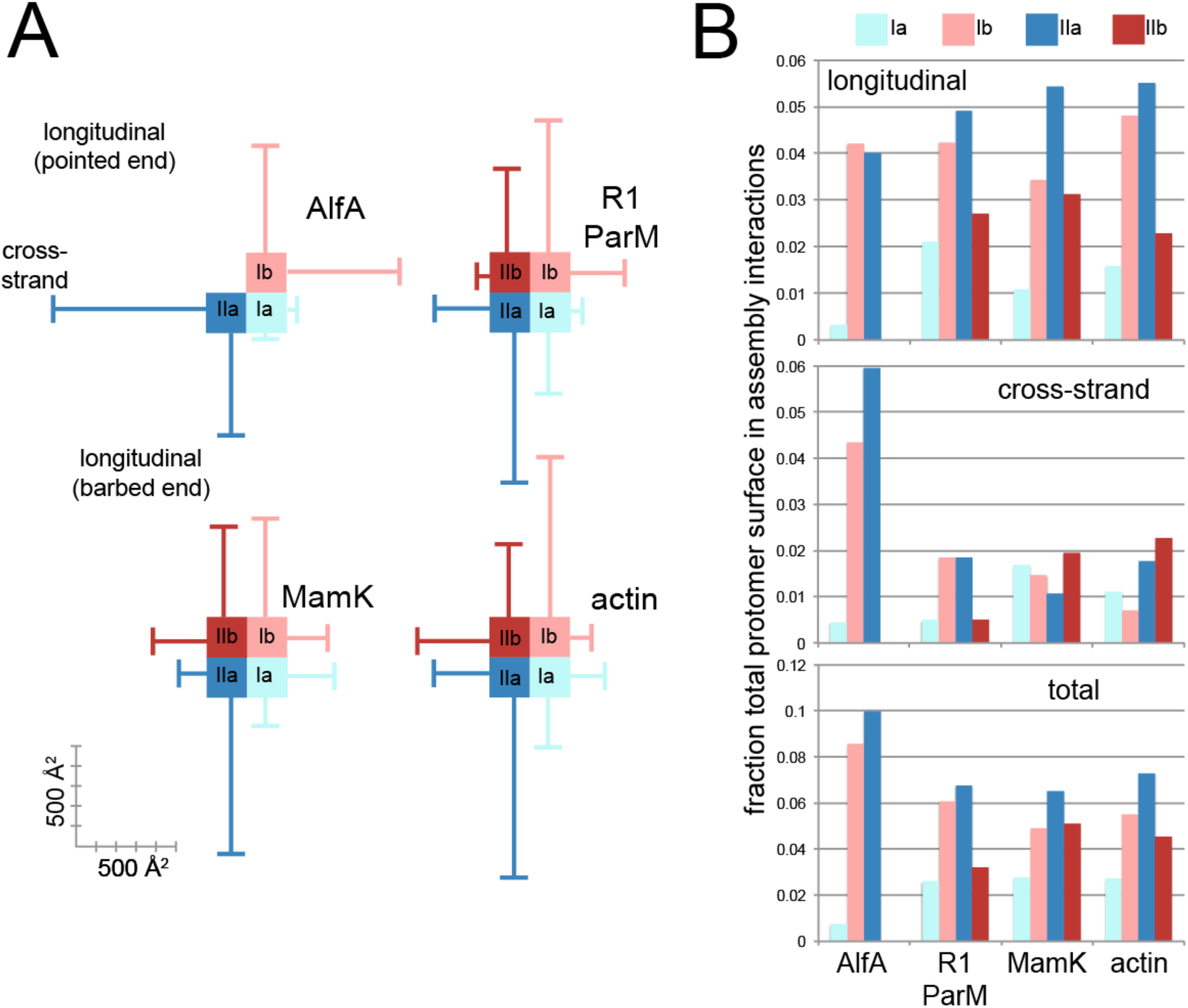
Evolutionary variation in the contribution of different subdomains to actin assembly interfaces. A) For each actin the area of interaction surfaces for each subdomain is plotted, showing contributions to longitudinal (vertical lines) and cross-strand (horizontal lines) interfaces. Scale bar at the bottom left indicates size of interfaces in Å^2^. B) The relative size of assembly interfaces for each subdomain are plotted as a fraction of total protomer surface area.

**Supplementary Figure 6.**
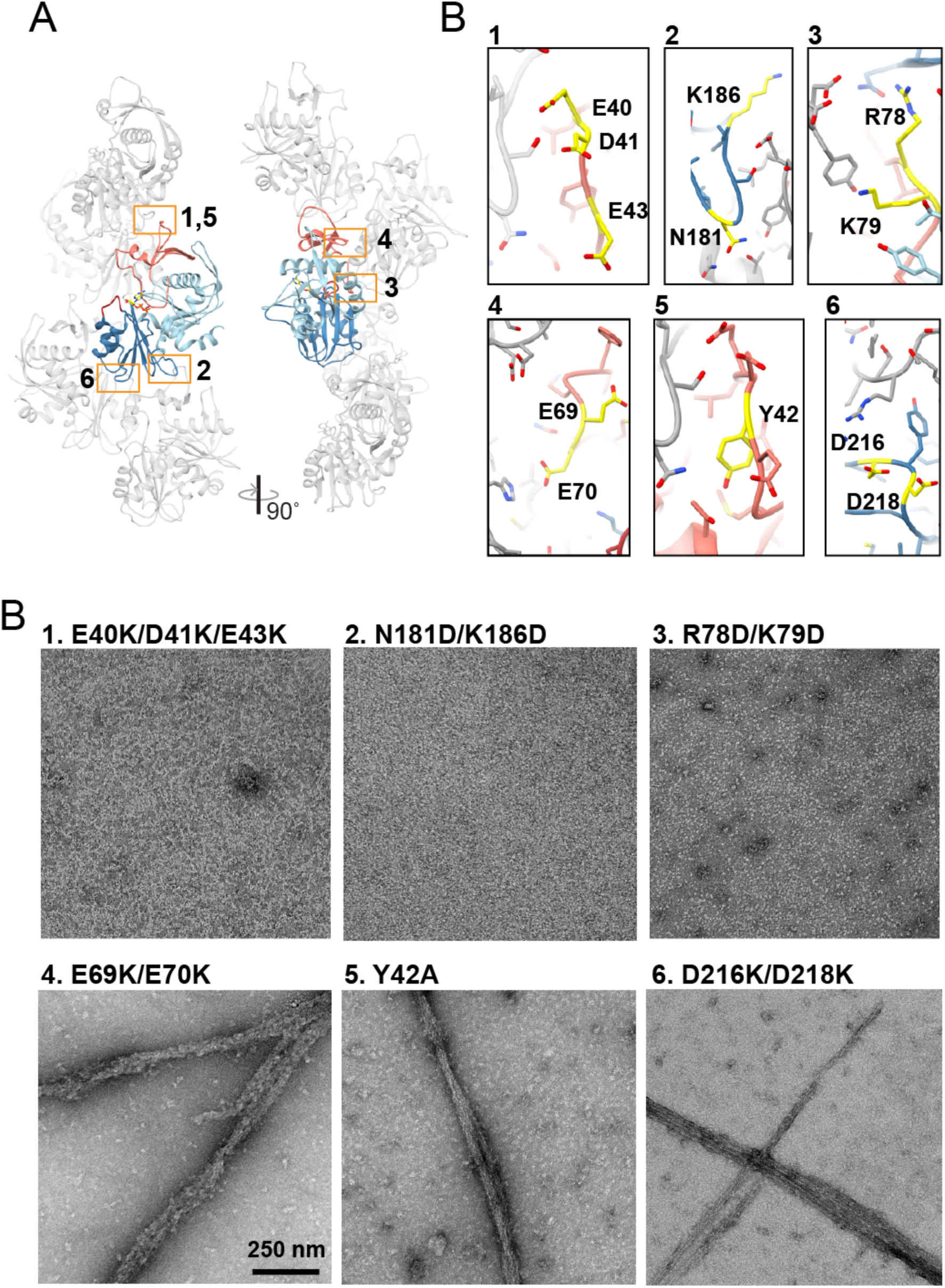
AlfA assembly mutants. Six mutants were designed on the basis of and AlfA homology model fit into a preliminary 12 Å reconstruction of the AlfA filament. A) Atomic model of the AlfA filament with a single protomer colored as in Figure 1, with locations of designed assembly mutants indicated by orange boxes. B) Close up views of the different designed mutations (yellow) in the final refined structure. C) Negative stain images of AlfA mutants showed that three designed mutations block assembly while three fail to prevent assembly. Inspection of the atomic model generated from high resolution cryo-EM structure (B) reveals that while mutants 4-6 are near assembly interfaces the sidechains of the mutated residues are not directly involved in contact, explaining why these mutants fail to prevent assembly.

